# Circular DNA vcDNA-S7 derived from segmented-dsRNA virus BmCPV attenuated viral infection through RNase H1

**DOI:** 10.1101/2024.04.17.590017

**Authors:** Jun Pan, Xinyu Tong, Yuming Ding, Chao Lei, Shulin Wei, An Xu, Yaxuan Han, Qunnan Qiu, Huilin Pang, Xiaolong Hu, Chengliang Gong, Min Zhu

## Abstract

Viral circDNA derived from insect positive-sense single-stranded RNA viruses plays a crucial role in regulating viral infections depending on the RNAi pathway. Our previous research has revealed that *Bombyx mori* cypovirus (BmCPV), a segmented-dsRNA virus, can generate viral circDNA vcDNA-S7, which can be transcribed to form RNA and further processed into small RNAs to defend against the virus through RNAi. However, whether vcDNA-S7 employs other mechanisms to resist the viruses. Presently, vcDNA-S7 has been found to resist BmCPV infection via a novel mechanism that depends on ribonuclease RNase H1 activity. It was found that the expression level of RNase H1 increased upon BmCPV infection (RNA virus) but not upon *B. mori* nucleopolyhedrovirus (BmNPV) (DNA virus) infection. RNase H1 has been shown to negatively regulate BmCPV infections. In addition, vcDNA-S7 was found to form a heteroduplex with the viral RNA and subsequently recruit RNase H1 to degrade the viral RNA within the heteroduplex, ultimately leading to the suppression of viral replication. Furthermore, the resistance of vcDNA-S7 to viruses was significantly reduced when cellular RNase H1 gene was silenced. These findings not only reveal a new antiviral mechanism involving circular DNA formed by reovirus but also offer intriguing insights into the antiviral function of RNase H1 in cells.

## 1 Introduction

Various mechanisms have been developed among bacteria and higher organisms to identify and eliminate alien genetic material, or to limit its functionality and replication. Nucleic acid-based defense systems, including Crisper Cas9 and RNAi, play an important role in antiviral processes [1, 2]. RNAi is an important pathway for counteracting infections caused by RNA viruses in insects [3, 4]. In the classic RNAi antiviral pathway, when the host encounters double-stranded RNAs (dsRNAs) from a virus’s genome or replication intermediate, the host’s Dicer-2 cuts these RNAs into smaller viral small interfering RNAs (vsiRNAs). The efficacy of the RNAi pathway in combating viruses is contingent upon the amplification of the RNAi effect, which typically relies on RNA-dependent RNA polymerase (RdRP) [5]. Recent studies have revealed the existence of a novel RNAi amplification mechanism in insects lacking RdRP [5]. For instance, once positive-sense single-stranded RNA viruses such as Flock house virus (FHV), *Drosophila* C virus (DCV), and Chikungunya virus (CHIKV) infects *Drosophila*, viral genomic RNAs can form stable viral circular DNAs that can be continuously and stably transcribed into a large amount of viral RNAs without relying on canonical transcriptional elements, thus producing highly abundant vsiRNAs to resist viral infection [5]. However, by knocking out Dicer, which is responsible for siRNA production, RNAi-deficient *Drosophila* could still resist virus infection mediated by viral DNAs [6], suggesting that there may be other antiviral mechanisms mediated by viral DNAs that are independent of the RNAi pathway in insects.

Previous studies have only involved non-retroviral single-stranded RNA viruses and only in a few insects, such as *Mosquitoes* and *Drosophila*. Further research is needed to ascertain the universality of the antiviral mechanism in insects, particularly their capacity to hinder viral infection by generating viral DNAs through the RNAi pathway. Silkworms, with their well-defined genetic background, serve as an excellent model insect for such studies. Our previous study showed that a typical species of reovirus, *B. mori* cypovirus (BmCPV), whose genome consists of 10 segmental double-stranded RNAs (S1-S10), could generate virus circular DNA vcDNA-S7 during infection in silkworm, and the RNAs transcribed by the vcDNA-S7 were found to be processed to vsiRNAs and further resist BmCPV infection by RNAi pathway, as in *Drosophila* [7], suggesting that the inhibition of viral infection through RNAi pathway mediated by viral circular DNAs is a common strategy in insects. However, another antiviral mechanism independent of the RNAi pathway based on virus circular DNAs in insects remains to be studied.

RNase H is a ubiquitous non-sequence-specific cellular endonuclease that is widely expressed and solely hydrolyzes the RNA of the RNA-DNA heteroduplex [8], which plays crucial roles in the biochemical processes associated with DNA replication, repair, and transcription where RNA-DNA hybrids occur. In addition, RNase H plays a crucial role in nucleic acid immunity by breaking down RNA-DNA hybrids that are formed during viral replication [9, 10]. It has been demonstrated in previous research that a lack of RNase H2, a member of the RNase H family, results in an accumulation of cytosolic RNA-DNA hybrids and an elevated level of retroelements, both of which can act as ligands for the cGAS–STING natural immune pathway [11, 12]. Recently, RNase H was found to be involved in a novel antiviral mechanism in mammalian cells based on nucleic acid immunity. It has been reported that negative-sense single-stranded virus complementary DNAs produced from the positive-sense single-stranded RNA virus encephalomyocarditis virus (EMCV) can form DNA-RNA hybrids with viral RNA, and DNA-RNA hybrids recruit RNase H to degrade the RNA strand of the DNA-RNA duplex, thus inhibiting EMCV infection [10]. Could circular DNA vcDNA derived from double-stranded RNA viruses defend against viral infection through RNase H1 in insects?

In this study, we examined whether vcDNA-S7, derived from segment S7 of reovirus BmCPV, could impede viral infection through RNase H1. Our findings reveal that the expression of RNase H1 increases during BmCPV infection and that RNase H1 negatively regulates the infection of the virus. Furthermore, vcDNA-S7 can combine with the corresponding viral RNA to form a heteroduplex, which then recruits RNase H1 to degrade the viral RNA in the heteroduplex. This process leads to the inhibition of viral proliferation. The findings indicate that vcDNA-S7 has the ability to obstruct BmCPV infection through RNase H1 involvement, uncovering a newly discovered nucleic acid-based antiviral mechanism in insects.

## 2 Results

### 2.1 Expression level of RNase H1 increased with BmCPV infection

RNase H1 has been reported to play a vital role in defense against single-stranded RNA virus infection [10]. We analyzed the expression levels of RNase H1 in BmCPV-infected and BmN cells at 0, 24, 48, 72, and 96 h post-infection. Our findings indicated that the RNase H1 expression level increased over time and reached its peak at 96 h post-infection (Fig 1A). This result was further confirmed by western blot analysis (Fig 1B and C). In order to directly observe the variation of RNase H1, we visualized the protein using immunofluorescence and evaluated its expression level by counting the number of infected cells that showed positive staining for RNase H1. We found that the proportion of red fluorescent cells, representing RNase H1 expression, increased with BmCPV infection (Fig 1D and E). RNA viruses have been shown to positively affect the expression levels of RNase H1. We also collected BmN cells infected with BmNPV, a DNA virus, at different infections. No significant changes in RNase H1 expression levels were observed in cells infected with BmNPV using real-time PCR (Figure 2A), western blotting (Fig 2B-C), and immunofluorescence (Fig 2D and E), suggesting that DNA virus didn’t affect the expression levels of RNase H1.

**Figure 1.**
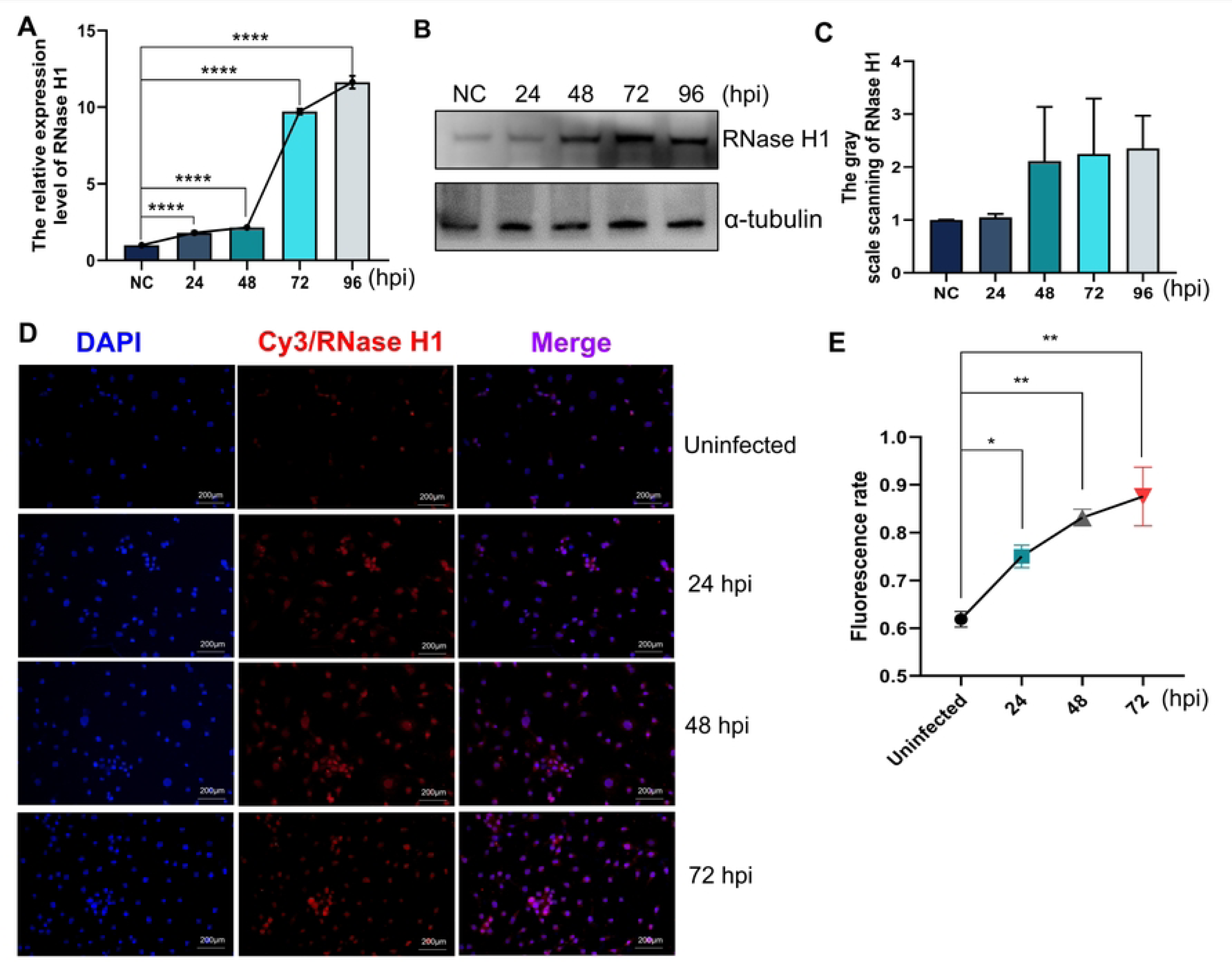
BmCPV infection altered the expression level of RNase H1. **A**, The express pattern of RNase H1 at different BmCPV infection periods by real-time PCR. **B**, The express pattern of RNase H1 was detected using Western blotting at different BmCPV infection periods. **C**, Gray value statistics for western blotting bands in panel B using the software Image J. **D**, Immunofluorescence assay were carried out using a rabbit anti-RNase H1 polyclonal antibody as the primary antibodies and using Cy3 (Red) conjugated goat anti-rabbit IgG(H+L) as the secondary antibodies. The cell nucleus was stained with DAPI (Blue). The cells were observed under a confocal microscope. **E,** Fluorescence statistics. All photographs taken were counted in the software Image J for the amount of blue/red fluorescence. The pictures were automatically adjusted in the software to highlight the contrast of the target, then, automatically counts the number of particles, and calculate the percentage of the red fluorescence numbers to the blue fluorescence numbers. (*, *P*≤0.05; **, *P*≤0.01; ***, *P*≤0.001, n=3)

**Figure 2.**
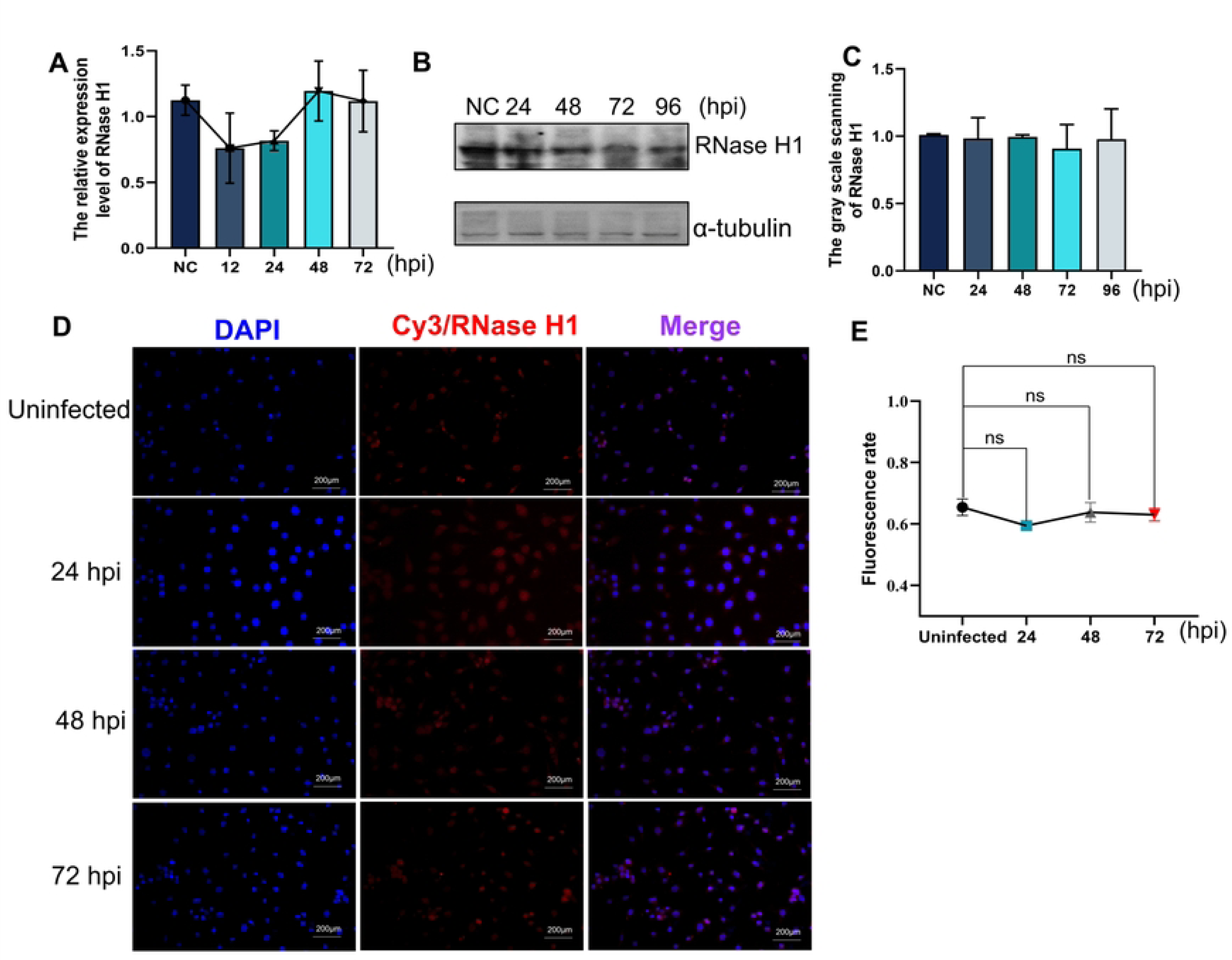
BmNPV infection cannot change the expression of RNase H1. **A**, The express pattern of RNase H1 at different BmNPV infection periods by real-time PCR. **B**, The express pattern of RNase H1 was detected using Western blotting at different BmCPV infection periods. **C**, Gray value statistics for western blotting bands in panel B using the software Image J. **D**, Immunofluorescence assay were carried out using anti-RNase H1 rabbit polyclonal antibody as the primary antibodies and using Cy3 (Red) conjugated goat anti-rabbit IgG(H+L) as the secondary antibodies. The cell nucleus was stained with DAPI (Blue). The cells were observed under a confocal microscope. **E,** Fluorescence statistics. The method is the same as in the above Figure 1E. (ns, no significance, n=3)

### 2.2 RNase H1 negatively regulates BmCPV proliferation

To explore the impact of RNase H1 on BmCPV proliferation, we detected changes in viral gene expression levels in cells with upregulated and downregulated RNase H1 expression. The results showed that when RNase H1 expression was silenced, mRNA level of BmCPV S1 increased (Fig 3A, C). Cells with overexpressed RNase H1 displayed contradictory results (Fig 3B, C). In addition, when RNase H1 expression was silenced, the expression level of VP7 protein also increased (Fig 3D-F), while overexpressing RNase H1, the level of VP7 decreased (Fig 3G-I), suggesting that RNase H1 negatively regulates BmCPV infection.

**Figure 3.**
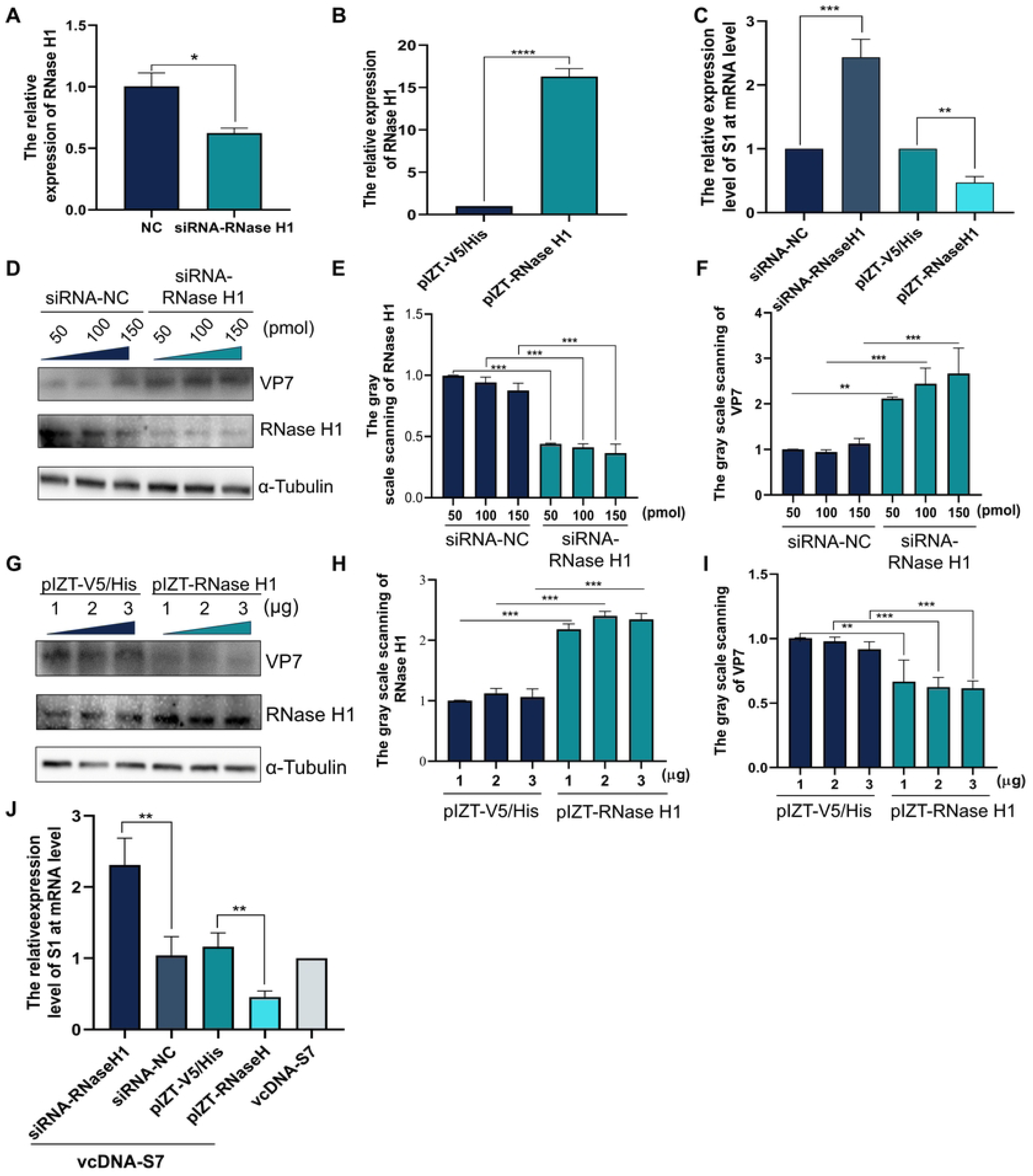
RNase H1 negatively regulates BmCPV infection. **A**, Silencing efficiency of siRNAs on RNase H1. BmN cells (1×10^6^) were transfected with RNase H1 siRNA for 48 h. Total RNA was extracted and the expression level of RNase H1 relative to TIF-4A was detected by real-time PCR. **B,** The mRNA level of RNase H1 were analyzed by real-time PCR in BmN cells transfected with pIZT-RNase H1. **C,** RNase H1 could negative regulate the mRNA level of BmCPV S1 gene to TIF-4A gene. 1×10^6^ BmN cells were pre-transfected with pIZT-RNase H1 or RNase H1 siRNA for 48 h, and total RNA was extracted and the expression level of S1 relative to TIF-4A was detected by real-time PCR. **D**, the protein level of VP7 and RNase H1 was analyzed by Western blotting when RNase H1 was silenced. BmN cells (1×10^6^) were transfected with RNase H1 siRNA with different dose (50 pmol, 100 pmol, 150 pmol) for 48 h, and then were infected with BmCPV for another 48 h. The total protein was extracted and the protein level of VP7 and RNase H1 was detected by Western blotting. The primary antibodies were VP7 mouse polyclonal antibody (diluted at 1:1000) and RNase H1 polyclonal antibody (diluted at 1:2000). The HRP-labeled goat anti-mouse IgG was used as the secondary antibody at 1:5000 dilution. **E** and **F,** Gray value statistics for western blotting bands in panel D using the software Image J. **G**, The protein level of VP7 and RNase H1 were analyzed by western blotting. 1×10^6^ BmN cells were transfected with different dose of pIZT-RNase H1 at 1 μg, 2 μg, 3 μg. The total protein was extracted and the protein level of VP7 and RNase H1 were detected by Western blotting. The VP7 mouse polyclonal antibody (diluted at 1:1000) and RNase H1 polyclonal antibody (diluted at 1:2000) were used as the primary antibodies. The HRP-labeled goat anti-mouse IgG was used as the secondary antibody at 1:5000 dilution. **H and I**, Gray value statistics for western blotting bands in panel F using the software Image J. **J,** VcDNA-S7 could regulate BmCPV infection through RNase H1. To explore whether vcDNA-S7 regulates BmCPV infection through RNase H1, the effect of vcDNA-S7 on BmCPV virus infection with overexpression of RNase H1 or knocking down of vcDNA-S7 was detected by real-time PCR. (*, *P*≤0.05; **, *P*≤ 0.01; ***, *P*≤0.001,n=3)

### 2.3 vcDNA-S7 regulates BmCPV infection through RNase H1

In our previous studies, vcDNA-S7 derived from BmCPV was discovered to be transcribed into RNA, which was then processed into vsiRNAs. This led to the suppression of BmCPV infection through the RNA interference pathway. Could vcDNA-S7 inhibit BmCPV infection via RNase H1? We examined the effect of RNase H1 expression on the ability of vcDNA-S7 to inhibit BmCPV infection. The results showed that when the expression of RNase H1 was increased, the ability of vcDNA-S7 to inhibit BmCPV was enhanced. Conversely, the ability of vcDNA-S7 to inhibit BmCPV was weakened when RNase H expression was reduced (Fig 3J). These results indicate that vcDNA-S7 could regulate BmCPV infection through RNase H1.

### 2.4 vcDNA-S7 exists in cells in two forms: single-stranded and double-stranded

Positive-sense single-stranded RNA virus-derived negative single-stranded circDNA could form a DNA-RNA duplex with viral RNA and recruit RNase H1 to degrade the RNA strand in the duplex structure, resulting in the inhibition of virus infection [10]. In our previous study, circular DNA vcDNA-S7 derived from BmCPV was found to inhibit BmCPV infection[7]. Can vcDNA-S7 form a DNA-RNA duplex with viral RNA and inhibit BmCPV infection through RNase H1? To study the novel antiviral mechanism of vcDNA-S7, it is crucial to determine whether vcDNA-S7A is a single-stranded or double-stranded circular DNA. It is widely acknowledged that DNA restriction endonucleases cleave double-stranded DNA (dsDNA) but no single-stranded DNA (ssDNA) [13]. Herein, the DNA extracted from the cells infected with BmCPV was digested by the restrictive endonuclease Sac II and used as a template for PCR to determine the form of vcDNA-S7 in nature using three pairs of primers (Primer1-F/R, Primer2-F/R, and Primer3-F/R) (Fig 4A). Figure 4 displays that the PCR products P1 amplified by Primer1-F/R and P2 amplified by Primer2-F/R were obtained from the digested pMD19T-vcDNA-S7-JS using Sac II and Primer1-F/R and Primer2-F/R primers, respectively. However, the PCR product P3 amplified by Primer3-F/R could not be obtained from the same plasmid using Primer3-F/R, suggesting that Sac II completely cleaved the double-stranded DNA plasmid pMD19T-vcDNA-S7-JS (Fig 4B). Total DNA extracted from BmCPV-infected and uninfected silkworm cells was digested with plasmid-Safe ATP-dependent DNase to remove linear DNA and then digested with Sac II, followed by PCR amplification using Primer3-F/R primers. The BmCPV-uninfected cell group was found to have no amplifiable PCR product when Primer3-F/R was used, regardless of whether Sac II cleavage was carried out. However, in the BmCPV-infected cell group, the corresponding PCR product could be detected by Primer3-F/R, regardless of whether Sac II cleavage was performed (Fig 4C), indicating that the vcDNA-S7 formed during BmCPV infection is resistant to cleavage by Sac II in its single-stranded DNA form.

**Figure 4.**
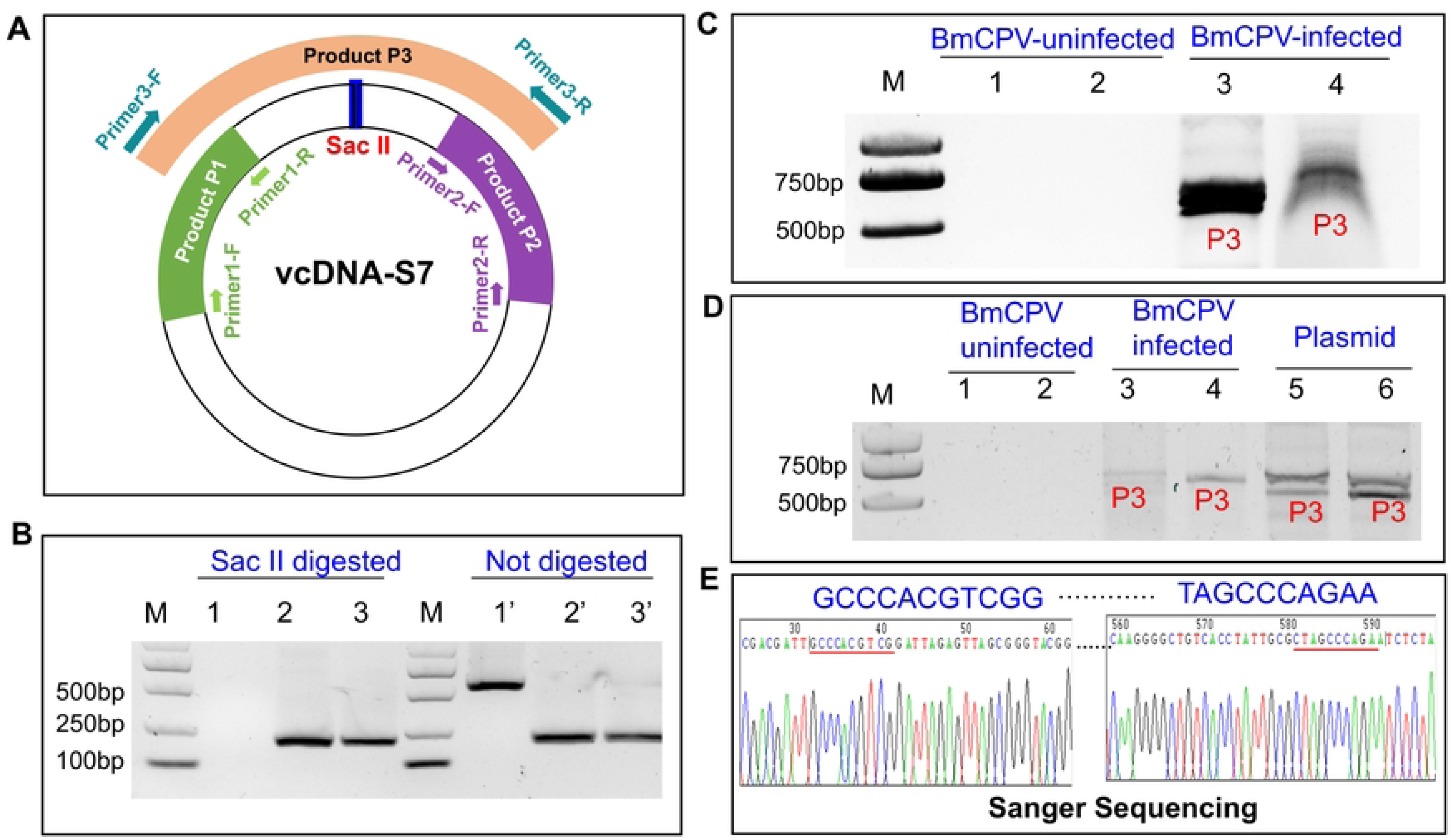
vcDNA-S7 exists in cells in two forms: single-stranded and double-stranded. **A**, Schematic diagram of the designed primers and the PCR products. **B**, Identification of pMD19T-vcDNA-S7-JS as a double stranded circular DNA. The adaptor sequence ligated with pMD19-T was digested with the restriction endonuclease Sac II. The digested production was identified by PCR with the designed primers. Lane M: DNA marker; Lane 1 and 1’: PCR products by Primer3-F/R; Lane 2 and 2’: PCR products by Primer1-F/R; Lane 3 and 3’: PCR products by Primer2-F/R. **C**, vcDNA-S7 has a double-stranded form. The genome extracted from BmCPV infected or un-infected BmN cells was digested by Sac II, and the integrity of DNA was evaluated by PCR with the primers Primer3-F/R. Lane M: DNA marker; Lane 1 and 3: PCR products using the DNA undigested with Sac II as template; Lane 2 and 4: PCR products using the DNA digested with *Sac*II as template. **D**, vcDNA-S7 has a single-stranded form. The genome extracted from BmCPV infected or un-infected BmN cells was incubated with 1 U nuclease S1. The integrity of DNA was evaluated by PCR with the primers Primer3-F/R. The recombinant plasmid pMD19T-vcDNA-S7-JS was used as a control. Lane M: DNA marker; Lane 1 and 3: PCR products using the DNA undigested with S1 as a template; Lane 2 and 4: PCR products using the DNA digested with S1 as template. Lane 5: PCR products using S1 undigested recombinant plasmid pMD19T-vcDNA-S7-JS as a template; Lane 6: PCR products using S1 digested recombinant plasmid pMD19T-vcDNA-S7-JS as a template. **E**, Sanger sequencing results of the PCR products of primer 3.

Moreover, after digestion with plasmid^TM^Safe ATP-dependent DNase, the DNA extracted from infected and uninfected BmCPV silkworms was further cleaved with nuclease S1 and PCR amplification was performed using Primer3-F/R primers. The nuclease-resistant pMD19T-vcDNA-S7-JS was subjected to the same treatment process. The results indicated that in the uninfected silkworm group, no Primer3-F/R product could be amplified after digestion with nuclease S1, while in the BmCPV-infected cell group, PCR products representing P3 could still be amplified by Primer3-F/R (Fig 4D), indicating that vcDNA-S7 also exists in a double-stranded form that is resistant to digestion with nuclease S1. Sanger sequencing results showed that the sequence of the PCR product amplified by Primer3-F/R from the infected cells was consistent with the theoretical sequence (Fig 4E). Collectively, the results indicated that vcDNA-S7 exists in infected cells in two forms: single-stranded and double-stranded, suggesting that vcDNA-S7 has the potential to form a DNA-RNA duplex with viral RNA.

### 2.5 DNA-RNA duplex formed by hybridization of vcDNA-S7 with BmCPV RNA recruits RNase H1

To confirm that vcDNA-S7 could formed a DNA-RNA duplex, an immunofluorescence co-localization assay was performed. The DNA-RNA duplex was labeled with red fluorescence (Cy3), and vcDNA-S7 was labeled with fluorescein isothiocyanate (FITC). The results showed that there was an overlap between green and red fluorescence in BmCPV infected BmN cells, but not in uninfected cells (Fig 5). Moreover, to determine the DNA-RNA duplex was formed by vcDNA-S7 and BmCPV S7 RNA in BmCPV-infected BmN cells, an immunofluorescence co-localization analysis of the complex was carried out. vcDNA-S7 was labeled with fluorescein isothiocyanate (FITC) represented by green. BmCPV S7 RNA was labeled with Cy5 represented by purple. and the DNA-RNA duplex was labeled with Cy3 represented by red. The results showed that there was an overlap between green, purple and red fluorescence in BmCPV infected BmN cells. However, the overlap and BmCPV S7 RNA significantly reduced after RNase H digestion, indicating that in BmCPV-infected cells, the DNA-RNA hybridization was formed by vcDNA-S7 and BmCPV S7 RNA, and RNase H1 could recognize the DNA-RNA hybridization to degrade BmCPV S7 RNA in the hybridization molecules (Fig 6).

**Figure 5.**
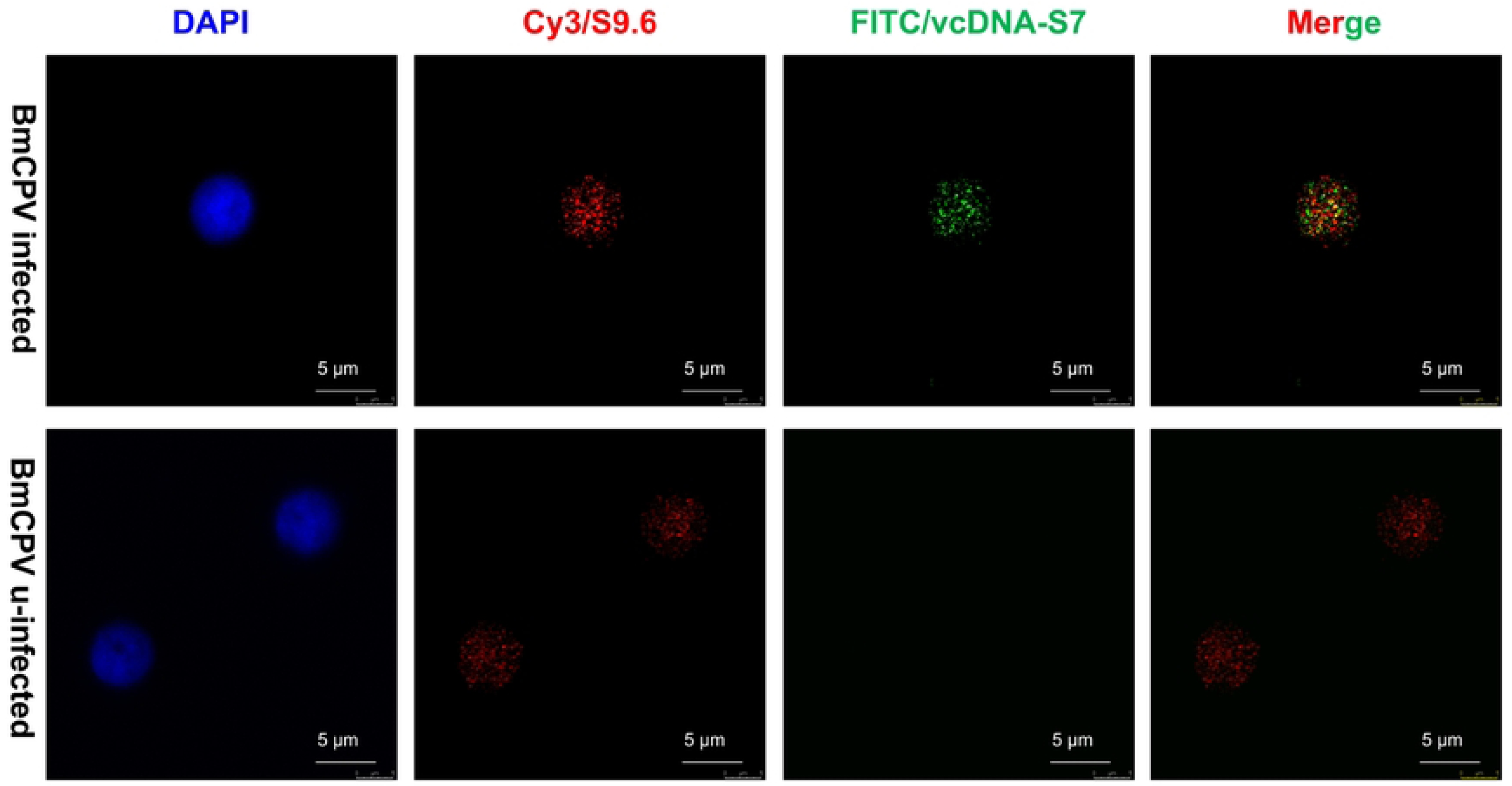
Co-immunofluorescence observation of vcDNA-S7 and DNA-RNA hybridization under BmCPV infection. Place cell climbing slices in a 24-well plate and inoculate with 10^4^ BmN cells. Add 5 μL of BmCPV polyhedra lysate (2×10^6^ polyhedra/mL) and infect for 48 hours. Subsequently, perform co-immunofluorescence experiments and observe under a confocal microscope. The primary antibodies used are S9.6 antibody (rabbit, 1:200) to detect DNA-RNA hybridization. vcDNA-S7 was detected by biotin-labeled probes. The secondary antibodies used are goat anti-mouse Cy3 antibody (1:200) and FITC-labeled streptavidin antibody. Cell nuclei are stained with DAPI solution (1:1000). DAPI represents labeled cell nuclei. FITC represents labeled RNase H1 protein. Cy3 represents labeled DNA: RNA hybrid chains, and Merge represents the combination of FITC and Cy3 fluorescence signals.

**Figure 6.**
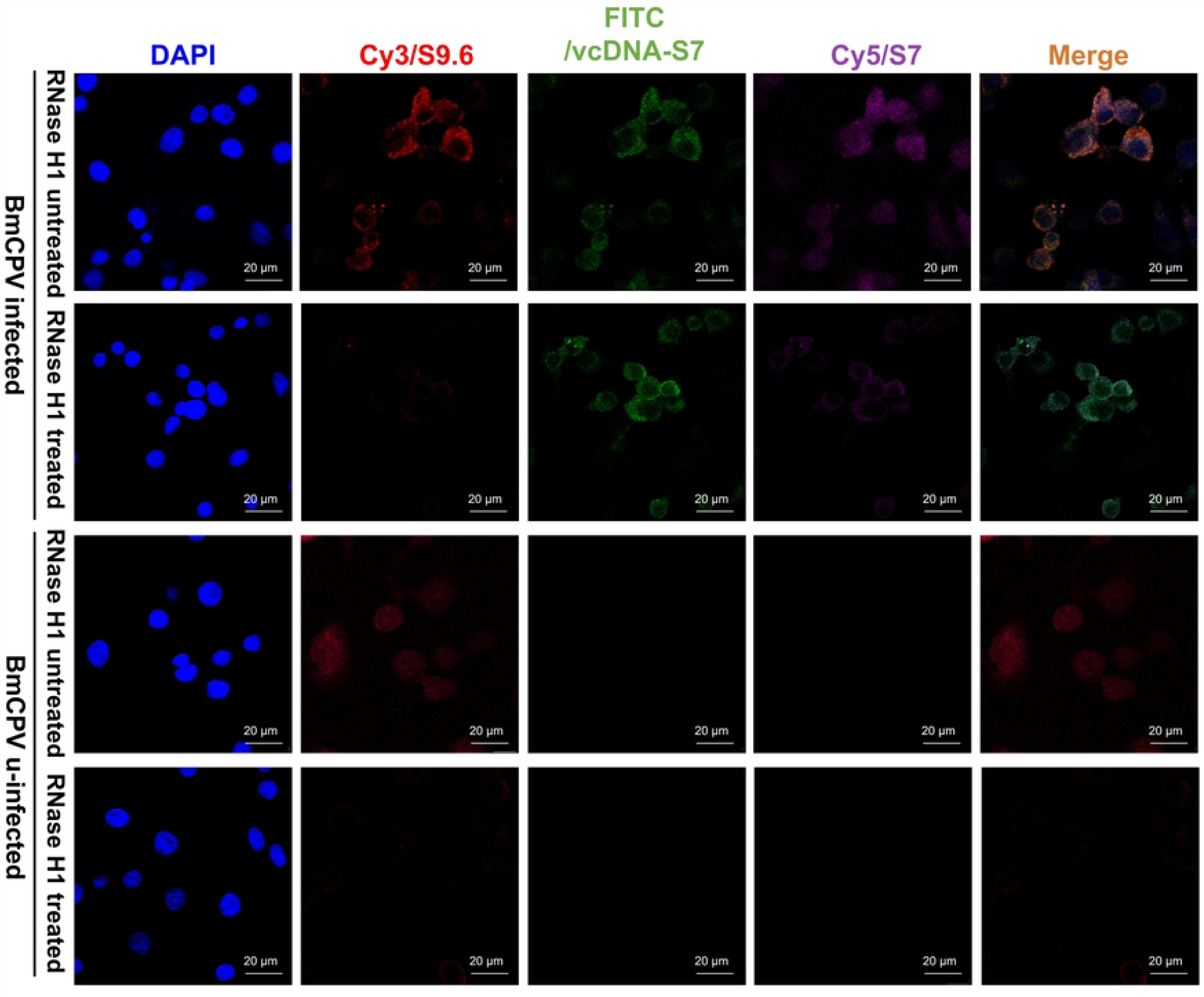
Co-immunofluorescence observation was performed on DNA-RNA hybridization formed by vcDNA-S7 and BmCPV S7 RNA recognized by RNase H1. BmN cells were seeded in a 24-well plate with 10^4^ cells per well and then infected with 5 μL of BmCPV polyhedra lysate (2×10^6^ polyhedra/mL). At 48 h post-infection, cells were collected and co-immunofluorescence experiments were conducted and observed under a confocal microscope. The primary antibodies used were S9.6 antibody (mouse, 1:200) to detect DNA-RNA hybridization, while the secondary antibodies used were goat anti-mouse Cy3 antibody (1:200). vcDNA-S7 and BmCPV S7 were detected by biotin-labeled probes and Cy5 probes, respectively. Cell nuclei were stained with DAPI (1:1000). DAPI represents labeled cell nuclei, Cy3 represents labeled DNA-RNA hybridization, FITC represents labeled vcDNA-S7, Cy5 represents labeled BmCPV S7 RNA and Merge represents the combination of FITC, Cy3 and Cy5 fluorescence signals.

Furthermore, a DNA-RNA immunoprecipitation (DRIP) experiment was conducted to validate DNA-RNA hybridization in BmCPV-infected cells (Fig 7A). The results revealed that the co-immunoprecipitation complex formed with the S9.6 antibody contained a greater amount of BmCPV S7 RNA and vcDNA-S7 than other samples. Following treatment with RNase H, the levels of enriched BmCPV S7 RNA and vcDNA-S7 in the immunoprecipitation complex decreased (Fig 7B). When using the RNase H antibody for immunoprecipitation, the results were the opposite (Fig 7C, D). These results indicated that vcDNA-S7 could form a DNA-RNA duplex with BmCPV S7 RNA and further recruit RNase H1 in the BmCPV-infected cells.

**Figure 7.**
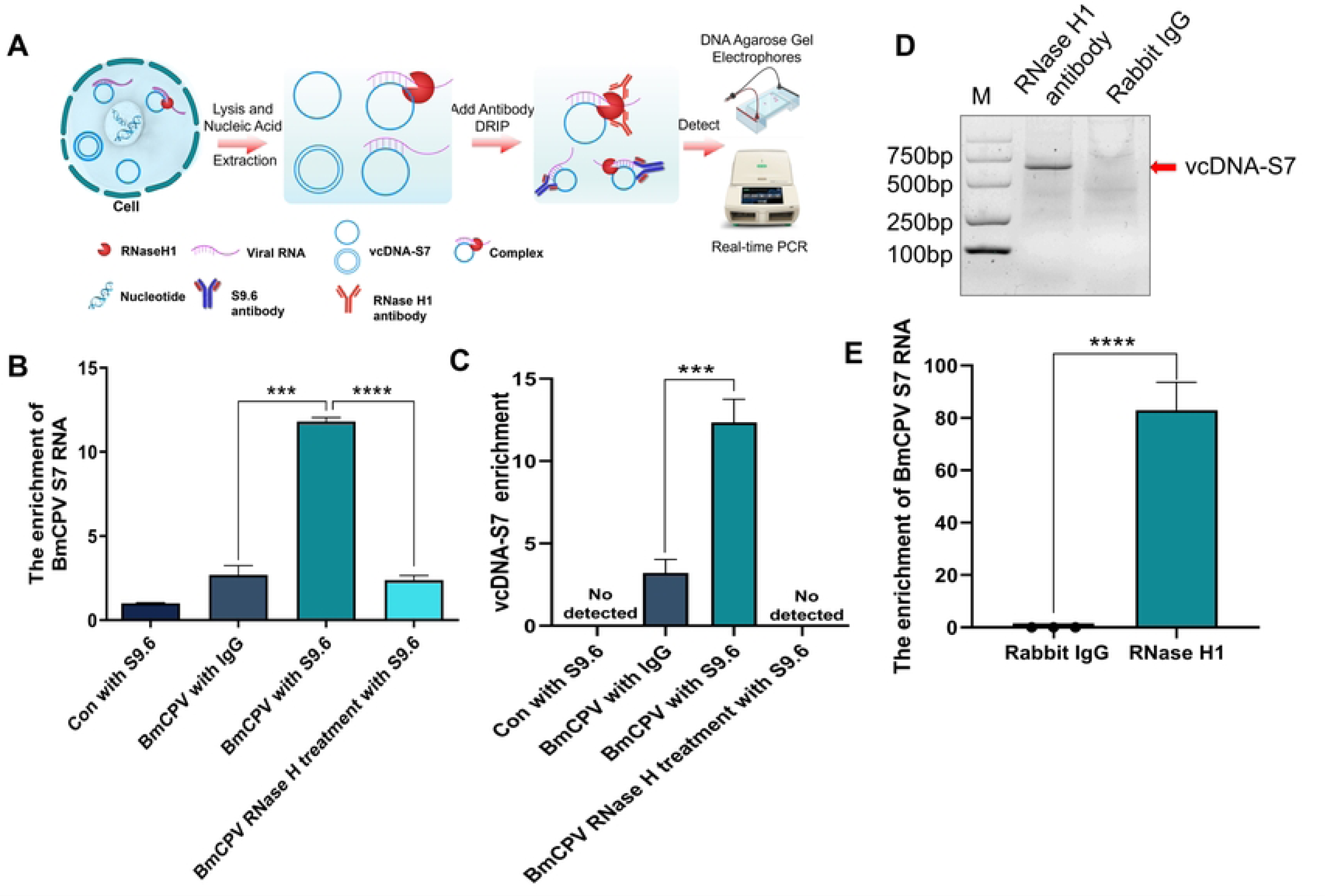
VcDNA-S7 could form DNA/RNA duplex with BmCPV RNA recruiting RNase H1. **A,** Experimental flowchart of DRIP. **B,** Immunoprecipitation assay based on S9.6 antibody for detection of BmCPV S7 RNA. 10^6^ BmN cells were inoculated with 10 μL of BmCPV polyhedra lysate (2×10^7^ polyhedra/mL). After 72 hours, nucleic acids were extracted and linear DNA was digested. 10% of the sample was taken as input, and the remaining samples were subjected to immunoprecipitation using IgG mouse and S9.6 antibodies, respectively. The immunoprecipitated complexes were enriched using Protein A+G agarose, and total RNA was extracted. BmCPV S7 RNA was detected using real-time PCR, and the ratio to the input was calculated. **C,** Immunoprecipitation assay based on S9.6 antibody for detection of vcDNA-S7. The immunoprecipitated complexes from part A were enriched using Protein A+G agarose, and nucleic acids were extracted. Real-time PCR was performed using specific primers to detect the relative copies of vcDNA-S7. Con with S9.6: Nucleic acid sample from uninfected cells, enriched with S9.6 antibody. BmCPV with IgG: Nucleic acid sample from BmCPV-infected cells, enriched with mouse IgG. BmCPV with S9.6: Nucleic acid sample from BmCPV-infected cells, enriched with S9.6 antibody. BmCPV RNase H treatment with S9.6: Nucleic acid sample from BmCPV-infected cells, treated with RNase H enzyme and enriched with S9.6 antibody. **D,** Immunoprecipitation assay based on RNase H1 antibody was used to detect vcDNA-S7. 10^6^ BmN cells were inoculated with 10 μL BmCPV polyhedra lysate (2×10^7^ polyhedra/mL). After 72 hours, nucleic acids were extracted and linear DNA was digested. Immunoprecipitation was performed using RNase H1 antibody, and the immunoprecipitated complex was enriched using Protein A+G agarose. Total DNA was extracted, and vcDNA-S7 was detected using specific primers through PCR. Meanwhile, A control group was set up using non-immune rabbit IgG for immunoprecipitation. **E,** Immunoprecipitation assay based on RNase H1 antibody was used to detect BmCPV S7 RNA. 10^6^ BmN cells were inoculated with 10 μL BmCPV polyhedra lysate (2×10^7^ polyhedra/mL). After 72 hours, nucleic acids were extracted and linear DNA was digested. 10% of the sample was taken as input, and the remaining sample was subjected to immunoprecipitation using RNase H1 antibody. The immunoprecipitated complex was enriched using Protein A+G agarose, and total RNA was extracted. BmCPV S7 RNA was quantified using real-time PCR, and the ratio to the input was calculated. (***, *P*≤0.001, n=3)

## 3 Discussion

RNase H is a non-sequence-specific endonuclease family that recognizes RNA-DNA hybrid molecules and catalyzes the degradation of RNA in RNA-DNA hybrid molecules through a hydrolysis mechanism. Based on evolution, the RNase H family can be broadly categorized into two subgroups, RNase H1 and RNase H2 [14]. Both RNase H1 and RNase H2 are involved in maintaining genomic stability, such as in the catalytic removal of loops [15–17]. In eukaryotes, RNase H1 participates in mitochondrial DNA replication [8, 18], and RNase H2 can perform ribonucleotide excision repair[19]. Retroviruses, such as human immunodeficiency virus (HIV), also have cellular RNase H1-like proteins. In these retroviruses, the RNase H domain exists in the reverse transcriptase proteins and is necessary for viral replication [20, 21]. In Hepatitis B virus (HBV), the RNase H domain is one of the four functional domains of HBV polymerase [22]. It is an indispensable enzyme for synthesizing negative-strand DNA of the HBV genome, which participates in the packaging of HBV DNA [20, 23]. Our studies suggest that RNase H is a crucial component of the viral life cycle. Specifically, we observed that the expression levels of silkworm RNase H1 are regulated by the RNA virus BmCPV, but not by the DNA virus BmNPV. This differential regulation of RNase H1 levels impacts the proliferation of BmCPV, but not BmNPV, implying that silkworm RNase H1 participates in the regulation of BmCPV infection.

Our previous study discovered that circular DNA called vcDNA-S7, derived from the RNA of BmCPV S7, can be transcribed into RNA and act as an antiviral agent through the RNAi pathway [7]. However, vcDNA-S7 employs other mechanisms to resist the viruses. Studies have shown that nucleic acid molecules can function by forming DNA-RNA hybrid molecules with homologous nucleic acid molecules through base pairing. In human breast cancer cells, the non-coding RNA circSMARCA5, produced by the chromatin remodeling factor 5 (SMARCA5) gene, can form DNA-RNA hybridization molecules with the DNA of its parent gene, which in turn regulates the transcription and translation of the SMARCA5 gene [24]. Recent studies have shown that, in mouse embryonic stem cells (mESCs), EMCV genomic RNA is reversibly transcribed to produce viral complementary DNAs under the mediation of cellular endogenous reverse transcriptase. Viral complementary DNAs can then form DNA-RNA hybrid molecules with viral RNAs. The intracellular endonuclease RNase H can specifically recognize DNA-RNA hybrid molecules and degrade viral RNA in the hybrid molecules, resulting in the inhibition of EMCV. This antiviral mechanism is known as endogenous transcriptase/RNase H-mediated antiviral system (ERASE) [10]. Based on the finding that vcDNA-S7 is involved in regulating BmCPV infection [7], this study further found that vcDNA-S7 can form a DNA-RNA hybrid molecule with BmCPV S7 RNA, and then recruit RNase H1 to degrade viral RNA in the DNA-RNA hybrid molecules to play an antiviral role. The antiviral mechanism of vcDNA-S7 is similar to that of viral complementary DNAs derived from EMCV in mouse mESCs. In insects, viral circular DNAs derived from positive-sense single-stranded RNA viruses have been reported to transcribe and produce vsiRNAs to inhibit the virus through the RNAi pathway [5, 25]. To date, ERASE has only been reported in positive-sense single-stranded RNA virus-infected mammalian cells [10]. Interestingly, vcDNA-S7 derived from the double-stranded RNA virus BmCPV can not only inhibit viral infection through RNAi, as in insects, but also through RNase H1, similar to mammalian cells. A previous study showed that viral circular DNAs are double-stranded in insects, which are formed through long terminal repeat (LTRs) mediation [5]. In mammalian cells, viral complementary DNAs exist in the form of single strands [10]. This study revealed that vcDNA-S7 exists in both single-and double-stranded forms and its ability to inhibit the virus may be attributed to its presence in both forms, via either the RNAi pathway or RNase H1.

Given that RNase H is prevalent in living organisms, has been conserved throughout evolution, and can be found in various organisms, including viruses, archaea, bacteria, and eukaryotes, it is important to take into account its presence and function [14]. Given the potential role of RNase H1 in regulating viral infection, we hypothesized that this enzyme may also play a similar role in other organisms. In other words, we speculated that endogenous reverse transcriptase-generating virus circular DNAs to suppress viral infection mediated by RNase H might be a common mechanism in various organisms as an anti-viral strategy. However, RNase H1 did not participate in host resistance to the DNA virus BmNPV. The ERASE mechanism is speculated to be a general strategy for host defense against RNA viruses, including double-stranded and single-stranded RNA viruses.

In summary, our study revealed that silkworm RNase H1 participates in the regulation of RNA virus BmCPV infection, but not DNA virus BmNPV. BmCPV-derived circular DNA vcDNA-S7 can form DNA/RNA hybridization molecules with BmCPV S7 RNA and recruit RNase H1 to degrade viral RNA in the hybridization molecules, thus defending against BmCPV infection (Fig 8). The findings of this study contribute a novel antiviral mechanism to the existing knowledge of silkworms. It has only been identified that eukaryotic mice and insects possess antiviral functions through reverse transcriptase and RNase H1. It will be interesting to study this antiviral mechanism in a broad range of organisms, including prokaryotic organisms.

**Figure 8.**
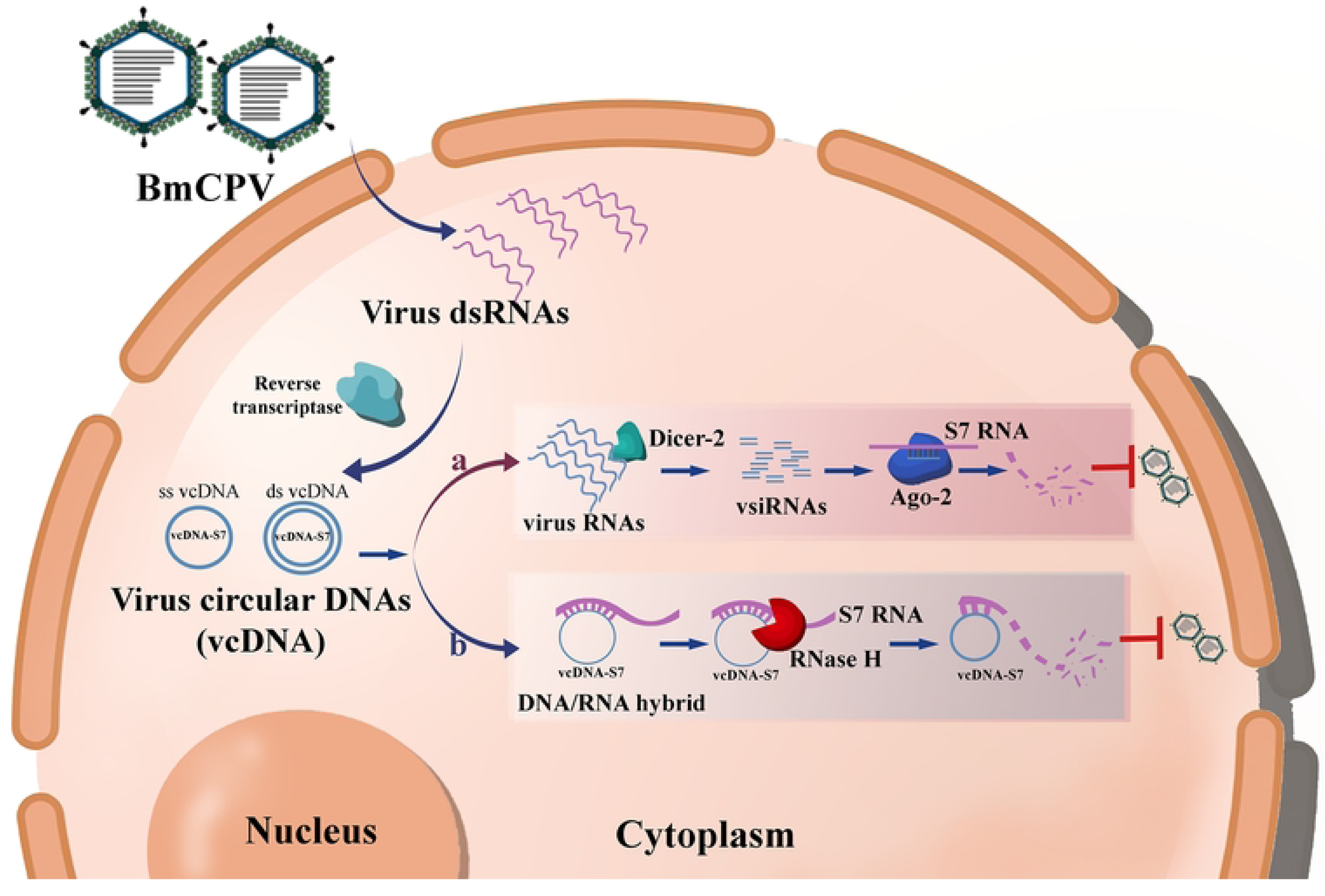
Circular DNA vcDNA-S7 derived from BmCPV attenuated virus infection mediated by RNase H1. BmCPV can form circular DNA vcDNA-S7 mediated by silkworm reverse transcriptase. (a), vcDNA-S7 can be transcribed to RNAs and produce vsiRNAs to control BmCPV infection by RNAi pathway. (b), vcDNA-S7 can form DNA/RNA hybridization molecules with BmCPV genome RNA and recruit RNase H1 to degrade viral RNA in the hybridization molecules, thus defending against BmCPV infection.

## 4 Experimental procedures

### 4.1 Cell culture

The BmN cells were cultured in TC-100 medium (Gibco, USA) supplemented with 10% fetal bovine serum (BI, Israel) at 26°C.

### 4.2 RNA extraction and reverse transcription

BmCPV-infected silkworm midgut or BmN cells were washed with pre-cooled 1×PBS. Total RNA was extracted using RNAiso Plus (Takara, Japan), according to the manufacturer’s instructions. The isolated RNAs were incubated with DNase I (Thermo Fisher Scientific, USA) to remove genomic DNA. RNA (4 μg of RNAs was reverse-transcribed using the EasyScript® First-Strand cDNA Synthesis SuperMix (TransGen Biotech, China) according to the manufacturer’s instructions.

### 4.3 DNA extraction and digestion

BmCPV-infected silkworm midguts or BmN cells were homogenized with pre-cooled 1×PBS. Thereafter, the DNA was purified according to the literature [10]. The isolated DNA was incubated with RNase A (Thermo Fisher, USA, Cat: EN0531) and plasmid-safe ATP-dependent DNase (Epicentre, USA, Cat: E3101K) mix at 37 °C for 30 min to remove RNAs and linear genomic DNA according to the instructions. Purified DNA was digested with Sac II (Takara, Japan) at 37℃ for 1 h. PCR was then performed using Sac II-digested and undigested DNA as templates. Three pairs of primers (Primer1-F/R, Primer2-F/R, and Primer3-F/R) were used to determine whether vcDNA-S7 was in a single- or double-stranded form. Primer1-F/R was designed upstream of the Sac II site, and primer Primer2-F/R was designed downstream of the Sac II site. Primer 3-F/R was designed to amplify the flanking sequence of Sac II (Fig 4A). The amplified fragment spanning the junction site of vcDNA-S7 was cloned into the pMD-19T vector (Takara, Dalian, China) using vcDNA-S7-specific primers to generate the positive control plasmid, pMD19T-vcDNA-S7-JS. Primer sequences are listed in Table S1.

### 4.4 Western blotting

Total protein (40 µg/lane) was separated by SDS-PAGE and transferred onto the PVDF membrane (Millipore, USA). After the membranes were blocked with 3% BSA in PBST containing 0.05% Tween-20, Western blotting was conducted with the respective primary antibodies. Anti-α-tubulin (Proteintech) was used as an internal reference. The secondary antibodies used were HRP-labeled goat anti-rabbit IgG or goat anti-mouse IgG at 1:5000 dilution (Proteintech, USA). Specific signal bands were visualized using an enhanced chemiluminescence (ECL) western blot detection kit (NCM Biotech, China). Quantitative analysis of the visible bands was performed using the ImageJ software.

### 4.5 Immunofluorescence

First, the cells or tissues were mounted onto coverslips or glass slides, followed by incubation with 1% bovine serum albumin (BSA) for 1 h at room temperature. Subsequently, the cells or tissues were incubated with a rabbit anti-RNase H1 antibody overnight at 4°C or for 1-2 hours at room temperature. Following the removal of unbound primary antibodies through washing with 0.1% Tween-20 in PBS, the cells or tissues were incubated with Cy3-labeled sheep anti-rabbit IgG for 1-2 h at room temperature. The cells or tissues were washed several times and stained with 4, 6-diamidino-2-phenylindole (DAPI, 1:1000). Finally, cell images were captured using a Leica DM2000 microscope (Leica, Germany). The ratio of red fluorescent cells was determined using the software Image J [26].

### 4.6 vcDNA-S7, BmCPV S7 RNA and DNA-RNA duplex co-location

BmN cells were seeded in 24-well plates at 10^4^/well and cultured for 24 h, after which they were infected with BmCPV. Cells were then collected at 48 h post-infection. After washing three times with 1× PBS, cells were fixed in 4% paraformaldehyde and permeabilized by 0.5% Triton X-100 for 20 min. According to previous study (Wu et al., 2021), the permeabilized coverslips were incubated with 5 U of RNase H (Abclonal, Wuhan, China) in 150 μl digestion buffer (20 mM Tris-HCl, pH 7.8, 40 mM KCl, 8 mM MgCl_2_, 1 mM DTT) for 1 h at room temperature for RNase H digestion. After two washes with PBS, cells were blocked with hybridization buffer for 4 h at 37 °C, followed by overnight incubation with cy5-labeled probes (targeting BmCPV S7 RNA) and biotin-labeled probes (targeting vcDNA-S7) in hybridization buffer (formula: 30 ml 20×SSC buffer (3M NaCl, 0.3M Na_3_Citrate·2H_2_O, adjust pH to 7.0), 10 ml 50×Denhardt’s, 5ml 10% SDS, 1ml 10 mg/ml Salmon DNA, 54 ml ddH_2_O). On the next day, 20×SSC buffer was diluted into 5×, 2×, 0.5× and 0.2×, and the slides were washed sequentially with 5×, 2×, 0.5× and 0.2× SSC buffer followed by blocked using blocking solution (3% BSA). Then, immunofluorescent staining was performed as above with corresponding antibodies. For immunofluorescence staining, cells were treated with the mouse S9.6 antibody (1:200), followed by fluorescein isothiocyanate (FITC)-conjugated anti-biotin antibodies (1:200) and CY3-conjugated goat anti-mouse IgG (1:200, Servicebio, GB21301, Wuhan, China). After washing, cells were counterstained with DAPI (1:1,000; Beyotime) and examined under Leica SP8 Confocal Microscope (Leica, Germany) with a 40× objective.

### 4.7 DNA-RNA immunoprecipitation (DRIP)

The nucleic acid was purified from the BmCPV-infected cells according to a previous study [10] and incubated with plasmid-safe ATP-dependent DNase I to remove linear genomic DNA. The DNase I-treated nucleic acid was dissolved in RNase/DNase-free water. The nucleic acid isolated from uninfected cells were used as a negative control. Ten percent of the DNase I-treated nucleic acid were used as input. The remaining nucleic acid samples were used for the detection of viral RNA or vcDNA-S7. Nucleic acid samples were treated with or without RNase H and then incubated with 5μg of S9.6 antibody (Abcam, USA) or mouse IgG (Beyotime, China) in the binding buffer (50mM Tris-HCl, pH 7.5, 150mM NaCl, 1mM EDTA, 1% Triton X-100, and 0.1% sodium deoxycholate) overnight at 4°C. The complex was incubated with Protein A+G (Beyotime, China) at 4 °C for 4 h. After being washed with PBS several times, the precipitate was dissolved in elution buffer (50mM Tris-HCl, pH 8.0, 10mM EDTA, 0.5% SDS). Then, the quantity of vcDNA-S7 and BmCPV S7 RNA was detected using real-time PCR with S7 divergent primers and S7 DNA primers (Table S1).

### 4.8 Immunoprecipitation with anti-RNase H1 antibody

BmN cells at 72 h post-infection with BmCPV were incubated with NP40 containing 1% PMSF to extract total cell protein. 400 μg of total cell protein was incubated with 5μL of anti-RNase H antibody (Abclonal, China) or rabbit IgG (Beyotime, China) at 4°C overnight. The complex was then incubated with Protein A+G (Beyotime, China) at 4 °C for 4 h. Following PBS washing several times, the precipitate was dissolved in the elution buffer for quantitative detection of vcDNA-S7 and BmCPV S7 RNA.

### 4.9 Construction of the plasmid pIZT-V5 /His-RNase H1

The gene RNase H1 (LOC101738648) was synthesized by Sangon Biotech Co., Ltd., and cloned into the pIZT-V5/His vector using restriction endonucleases Sac I and Sac II to construct the pIZT-V5/His-RNase H1 plasmid, which is capable of expressing the RNase H1 protein fused with a V5 tag.

### 4.10 RNAi

RNase H1-specific siRNAs were designed and synthesized by Sangon Biotech (Shanghai, China). A total of 100 pmol siRNA was transfected into 1×10^6^ BmN cells using Roch-X gem (Cat:6366236001; Roche, Switzerland). siRNA-NC was used as the negative control. At 48 h post-transfection, the cells were infected with BmCPV for an additional 48 h, followed by real-time PCR to evaluate the expression level of RNase H1. The siRNA sequences are listed in Table S2.

### 4.11 Quantitative Real-time PCR

To determine the expression level of the target gene, 1×10^6^ BmN cells were inoculated with BmCPV. Total RNA was extracted from infected cells and reverse-transcribed to cDNA using random primers (TransGen Biotech, China). The relative mRNA levels of the target genes were detected by real-time PCR using the primers listed in Table S1. The housekeeping gene *TIF-4A* was used as the normalizer.

### 4.12 Statistics

Data are expressed as means ± standard deviation (SD). Statistical analyses were performed by one-way analysis of variance (ANOVA) and t-test to determine statistical significance between the groups using GraphPad Prism 8 software. *P*-value significance was set at *P*≤0.05.

## Founding

This study was funded by the National Natural Science Foundation of China (32202744, 32372946 and 32072792), the Suzhou Agricultural Science and Technology Innovation Project (SNG2021033), the Natural Science Foundation of the Jiangsu Higher Education Institutions of China (22KJB23003), and Priority Academic Program of Development of Jiangsu Higher Education Institutions.

## Ethics approval and consent to participate

Not applicable

## Consent for publication

Not applicable

## Availability of data and materials

The data generated or used during the study are available from the corresponding author by reasonable request.

## Authors’ contributions

M.Z., and C.G. conceived and designed the experiments. J.P., X.T., Y.D., C.L., S.W., and A.X.performed the experiments. Y. H., Q.Q., H.P., and X.H. analyzed the experimental data. M.Z., and C.G. wrote and edited the manuscript.

## Acknowledgements

We thank for EditSprings (https://www.editsprings.cn) for the expert linguistic services provided.

---

**Table S1.**
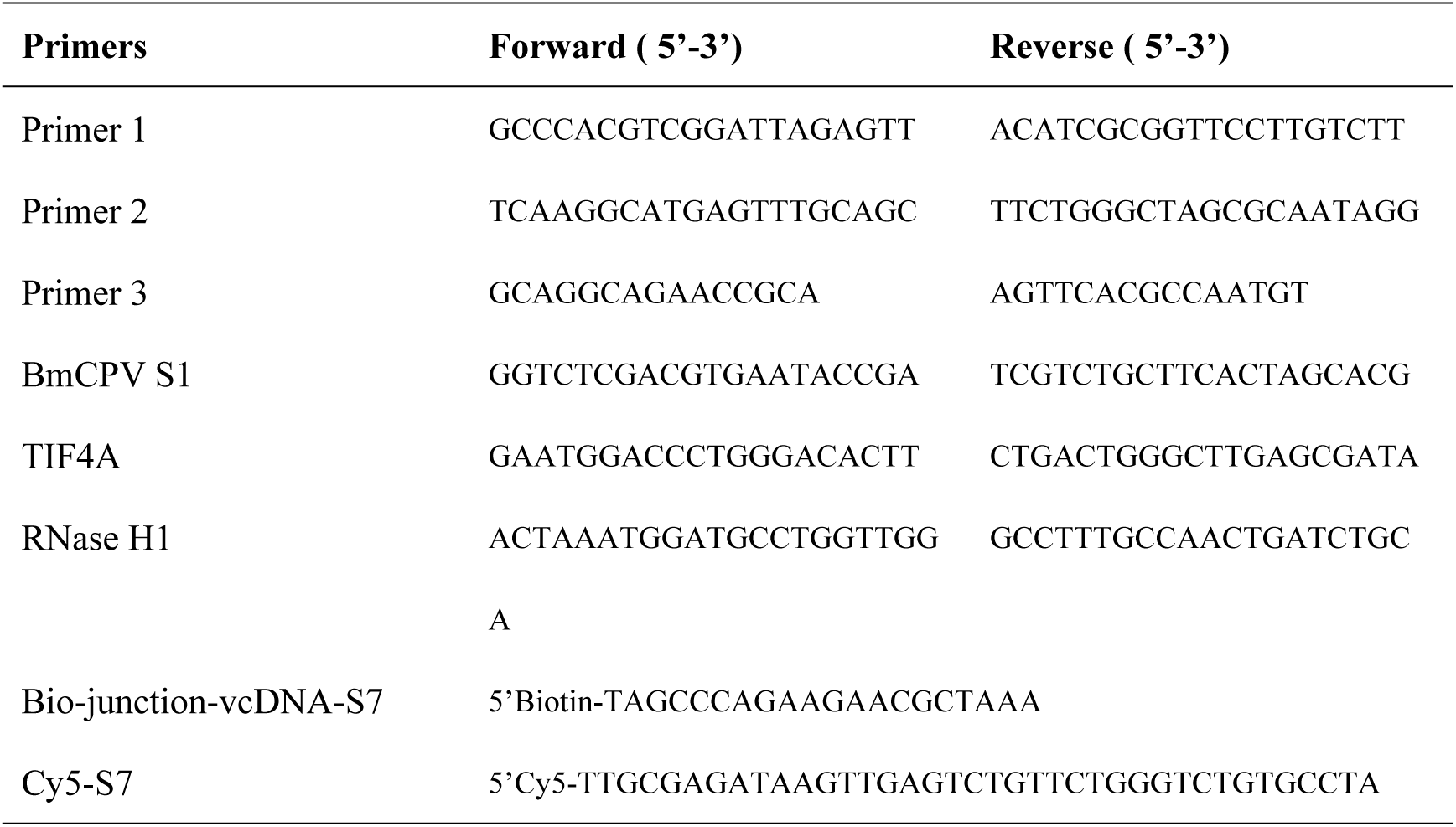
The sequences of primers and probes used in this study.

**Table S2.**
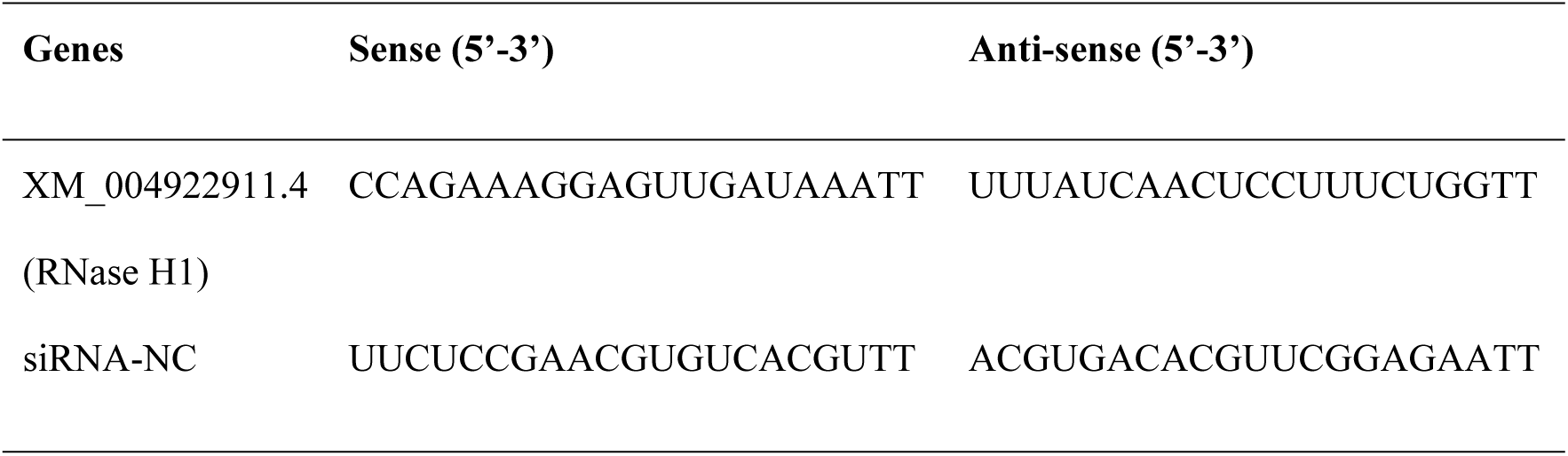
The siRNA sequencing used in this study.

## References

1. Jiang FG, Doudna JA. CRISPR-Cas9 Structures and Mechanisms. Annu Rev Biophys. 2017;46:505–29. doi: 10.1146/annurev-biophys-062215-010822. PubMed PMID: WOS:000402908700023.

2. Schuster S, Miesen P, van Rij RP. Antiviral RNAi in Insects and Mammals: Parallels and Differences. Viruses-Basel. 2019;11(5). doi: ARTN 448 10.3390/v11050448. PubMed PMID: WOS:000472676600059.

3. Zhu KY, Palli SR. Mechanisms, Applications, and Challenges of Insect RNA Interference. Annu Rev Entomol. 2020;65:293–311. doi: 10.1146/annurev-ento-011019-025224. PubMed PMID: WOS:000507471600015.

4. Jin Y, Zhao JH, Guo HS. Recent advances in understanding plant antiviral RNAi and viral suppressors of RNAi. Curr Opin Virol. 2021;46:65–72. doi: 10.1016/j.coviro.2020.12.001. PubMed PMID: WOS:000629333100009.

5. Poirier EZ, Goic B, Tomé-Poderti L, Frangeul L, Boussier J, Gausson V, et al. Dicer-2-Dependent Generation of Viral DNA from Defective Genomes of RNA Viruses Modulates Antiviral Immunity in Insects. Cell Host Microbe. 2018;23(3):353-+. doi: 10.1016/j.chom.2018.02.001. PubMed PMID: WOS:000427477400011.

6. Mondotte JA, Gausson V, Frangeul L, Blanc H, Lambrechts L, Saleh MC. Immune priming and clearance of orally acquired RNA viruses in. Nat Microbiol. 2018;3(12):1394–403. doi: 10.1038/s41564-018-0265-9. PubMed PMID: WOS:000451259600013.

7. Zhu M, Pan J, Tong XY, Qiu QN, Zhang X, Zhang YX, et al. BmCPV-Derived Circular DNA vcDNA-S7 Mediated by Reverse Transcriptase (RT) Regulates BmCPV Infection. Front Immunol. 2022;13. doi: ARTN 861007 10.3389/fimmu.2022.861007. PubMed PMID: WOS:000777473300001.

8. Cerritelli SM, Frolova EG, Feng CG, Grinberg A, Love PE, Crouch RJ. Failure to produce mitochondrial DNA results in embryonic lethality in null mice. Mol Cell. 2003;11(3):807–15. doi: Doi 10.1016/S1097-2765(03)00088-1. PubMed PMID: WOS:000182049700024.

9. Moelling K, Broecker F, Russo G, Sunagawa S. RNase H As Gene Modifier, Driver of Evolution and Antiviral Defense. Front Microbiol. 2017;8. doi: ARTN 1745 10.3389/fmicb.2017.01745. PubMed PMID: WOS:000410643100001.

10. Wu JY, Wu CY, Xing F, Cao L, Zeng WJ, Guo LP, et al. Endogenous reverse transcriptase and RNase H-mediated antiviral mechanism in embryonic stem cells. Cell Res. 2021;31(9):998–1010. doi: 10.1038/s41422-021-00524-7. PubMed PMID: WOS:000665637200001.

11. Mackenzie KJ, Carroll P, Lettice L, Tarnauskaite Z, Reddy K, Dix F, et al. Ribonuclease H2 mutations induce a cGAS/STING-dependent innate immune response. Embo J. 2016;35(8):831–44. doi: DOI 10.15252/embj.201593339. PubMed PMID: WOS:000374846600007.

12. Pokatayev V, Hasin N, Chon H, Cerritelli SM, Sakhuja K, Ward JM, et al. RNase H2 catalytic core Aicardi-Goutieres syndrome-related mutant invokes cGAS-STING innate immune-sensing pathway in mice. J Exp Med. 2016;213(3):329–36. doi: 10.1084/jem.20151464. PubMed PMID: WOS:000373390300004.

13. Smith DR. Restriction Endonuclease Digestion of DNA. In: Murphy D, Carter DA, editors. Transgenesis Techniques: Principles and Protocols. Totowa, NJ: Humana Press; 1993. p. 427–31.

14. Hyjek M, Figiel M, Nowotny M. RNases H: Structure and mechanism. DNA Repair. 2019;84. doi: ARTN 102672 10.1016/j.dnarep.2019.102672. PubMed PMID: WOS:000504779700008.

15. Broccoli S, Rallu F, Sanscartier P, Cerritelli SM, Crouch RJ, Drolet M. Effects of RNA polymerase modifications on transcription-induced negative supercoiling and associated R-loop formation. Mol Microbiol. 2004;52(6):1769–79. doi: 10.1111/j.1365-2958.2004.04092.x. PubMed PMID: WOS:000221866300019.

16. Chon H, Sparks JL, Rychlik M, Nowotny M, Burgers PM, Crouch RJ, et al. RNase H2 roles in genome integrity revealed by unlinking its activities. Nucleic Acids Res. 2013;41(5):3130–43. doi: 10.1093/nar/gkt027. PubMed PMID: WOS:000318062600035.

17. Lockhart A, Pires VB, Bento F, Kellner V, Luke-Glaser S, Yakoub G, et al. RNase H1 and H2 Are Differentially Regulated to Process RNA-DNA Hybrids. Cell Rep. 2019;29(9):2890-+. doi: 10.1016/j.celrep.2019.10.108. PubMed PMID: WOS:000499377000024.

18. Lima WF, Murray HM, Damle SS, Hart CE, Hung G, De Hoyos CL, et al. Viable knockout mice show RNaseH1 is essential for R loop processing, mitochondrial and liver function. Nucleic Acids Res. 2016;44(11):5299–312. doi: 10.1093/nar/gkw350. PubMed PMID: WOS:000379753100036.

19. Williams JS, Lujan SA, Kunkel TA. Processing ribonucleotides incorporated during eukaryotic DNA replication. Nat Rev Mol Cell Bio. 2016;17(6):350–63. doi: 10.1038/nrm.2016.37. PubMed PMID: WOS:000376526400010.

20. Tramontano E, Corona A, Menéndez-Arias L. Ribonuclease H, an unexploited target for antiviral intervention against HIV and hepatitis B virus. Antivir Res. 2019;171. doi: ARTN 104613 10.1016/j.antiviral.2019.104613. PubMed PMID: WOS:000499937000013.

21. Ilina TV, Brosenitsch T, Sluis-Cremer N, Ishima R. Retroviral RNase H: Structure, mechanism, and inhibition. Enzymes. 2021;50:227–47. Epub 20210924. doi: 10.1016/bs.enz.2021.07.007. PubMed PMID: 34861939; PubMed Central PMCID: PMCPMC8994160.

22. Clark DN, Tajwar R, Hu J, Tavis JE. The hepatitis B virus polymerase. Enzymes. 2021;50:195–226. Epub 20210809. doi: 10.1016/bs.enz.2021.06.010. PubMed PMID: 34861937.

23. Chen Y, Robinson WS, Marion PL. Selected Mutations of the Duck Hepatitis-B Virus-P Gene Rnase-H Domain Affect Both Rna Packaging and Priming of Minus-Strand DNA-Synthesis. J Virol. 1994;68(8):5232–8. doi: Doi 10.1128/Jvi.68.8.5232-5238.1994. PubMed PMID: WOS:A1994NW97800058.

24. Xu XL, Zhang JW, Tian YH, Gao Y, Dong X, Chen WB, et al. CircRNA inhibits DNA damage repair by interacting with host gene. Mol Cancer. 2020;19(1). doi: ARTN 128 10.1186/s12943-020-01246-x. PubMed PMID: WOS:000566695900002.

25. Goic B, Vodovar N, Mondotte JA, Monot C, Frangeul L, Blanc H, et al. RNA-mediated interference and reverse transcription control the persistence of RNA viruses in the insect model. Nat Immunol. 2013;14(4):396–403. doi: 10.1038/ni.2542. PubMed PMID: WOS:000316648700015.

26. Collins CM, Dunleavy EM. Imaging and Quantitation of Assembly Dynamics of the Centromeric Histone H3 Variant CENP-A in Drosophila melanogaster Spermatocytes by Immunofluorescence and Fluorescence In-Situ Hybridization (Immuno-FISH). Methods Mol Biol. 2018;1832:327–37. doi: 10.1007/978-1-4939-8663-7_18. PubMed PMID: 30073536.

